# The embryonic transcriptome of *Arabidopsis thaliana*

**DOI:** 10.1101/479584

**Authors:** Falko Hofmann, Michael A. Schon, Michael D. Nodine

## Abstract

Cellular differentiation is associated with changes in transcript populations. Accurate quantification of transcriptomes during development can thus provide global insights into differentiation processes including the fundamental specification and differentiation events operating during plant embryogenesis. However, multiple technical challenges have limited the ability to obtain high quality early embryonic transcriptomes, namely the low amount of RNA obtainable and contamination from surrounding endosperm and seed-coat tissues. We compared the performance of three low-input mRNA sequencing (mRNA-seq) library preparation kits on 0.1 to 5 nanograms (ng) of total RNA isolated from *Arabidopsis thaliana* (Arabidopsis) embryos and identified a low-cost method with superior performance. This mRNA-seq method was then used to profile the transcriptomes of Arabidopsis embryos across eight developmental stages. By comprehensively comparing embryonic and post-embryonic transcriptomes, we found that embryonic transcriptomes do not resemble any other plant tissue we analyzed. Moreover, transcriptome clustering analyses revealed the presence of four distinct phases of embryogenesis which are enriched in specific biological processes. We also compared zygotic embryo transcriptomes with publicly available somatic embryo transcriptomes. Strikingly, we found little resemblance between zygotic embryos and somatic embryos derived from late-staged zygotic embryos suggesting that the molecular basis of somatic and zygotic embryogenesis are distinct from each other. In addition to the biological insights gained from our systematic characterization of the Arabidopsis embryonic transcriptome, we provide a data-rich resource for the community to explore.

**Key Message:** Arabidopsis embryos possess unique transcriptomes relative to other plant tissues including somatic embryos, and can be partitioned into four transcriptional phases with characteristic biological processes.

## Introduction

Flowering plants begin their life as an embryo deeply embedded within a seed. In *Arabidopsis thaliana* (Arabidopsis), a series of stereotypical cell divisions produces the fundamental body plan that already possesses precursors to the shoot and root meristem, as well as the epidermal, ground and vascular tissues arranged in concentric layers. Cellular differentiation includes, and is largely defined by, changes in the quantity of specific transcripts present in the cell. Therefore, understanding cellular differentiation during embryogenesis requires the ability to quantify embryonic transcriptomes. However, multiple technical challenges limit the ability to obtain high quality embryo transcriptomes especially from the earliest stages when basic patterning processes are instrumental in defining the plant body plan. Due to the small size of early embryos and their enclosure within a seed, isolating RNA from embryos is prone to contamination from the surrounding endosperm and maternal sporophytic RNA (Schon and Nodine 2017). Additionally, the protocol must be sensitive enough to detect even lowly abundant transcripts from the few nanograms or less of total input RNA that is typically obtainable from early embryos.

## Results

### Comparison of low-input mRNA-seq library preparation methods

A variety of low-input mRNA sequencing (mRNA-seq) methods have been developed for tissue-specific and single-cell sequencing (reviewed in (Chen et al. 2018)). To determine the optimal mRNA-seq method for profiling transcriptomes from low-input total RNA isolated from Arabidopsis embryos, we compared the performance of three different mRNA-seq library construction protocols. We prepared mRNA-seq libraries from 5, 1, 0.5 or 0.1 ng of total RNA isolated from bent-cotyledon staged embryos using either the Ovation PicoSL WTA System V2 (Ovation; Nugen) or SMARTer Ultra Low Input RNA Kit for Sequencing - v3 (SMARTer; Clontech) commercially available kits, or the non-commercial Smart-seq2 method (Picelli et al. 2013).

We were able to detect between 13,453-16,315 protein-coding genes with at least 1 transcript per million (TPM) from libraries constructed with Smart-seq2 across the dilution series of input RNA (Fig. 1A). This is comparable to Ovation, which detected between 14,398-16,545 genes across the same range of RNA input. In contrast, libraries generated with SMARTer had lower sensitivity compared with the other two methods and only detected 6,218 unique protein-coding genes from 100 picograms of total RNA. Methods that enable the sequencing of full-length transcripts provide a more accurate representation of the transcriptome. We therefore determined which method captured full-length transcript sequences by comparing the coverage of mRNA-seq reads along Araport11-annotated protein-coding genes (Cheng et al. 2017) (Fig. 1B). mRNA-seq libraries generated with Smart-seq2 produced the most uniform coverage along protein-coding genes, while SMARTer library mRNA-seq reads were most abundant in the middle of transcripts and Ovation library reads showed uneven coverage that varied from gene to gene.

**Figure 1.**
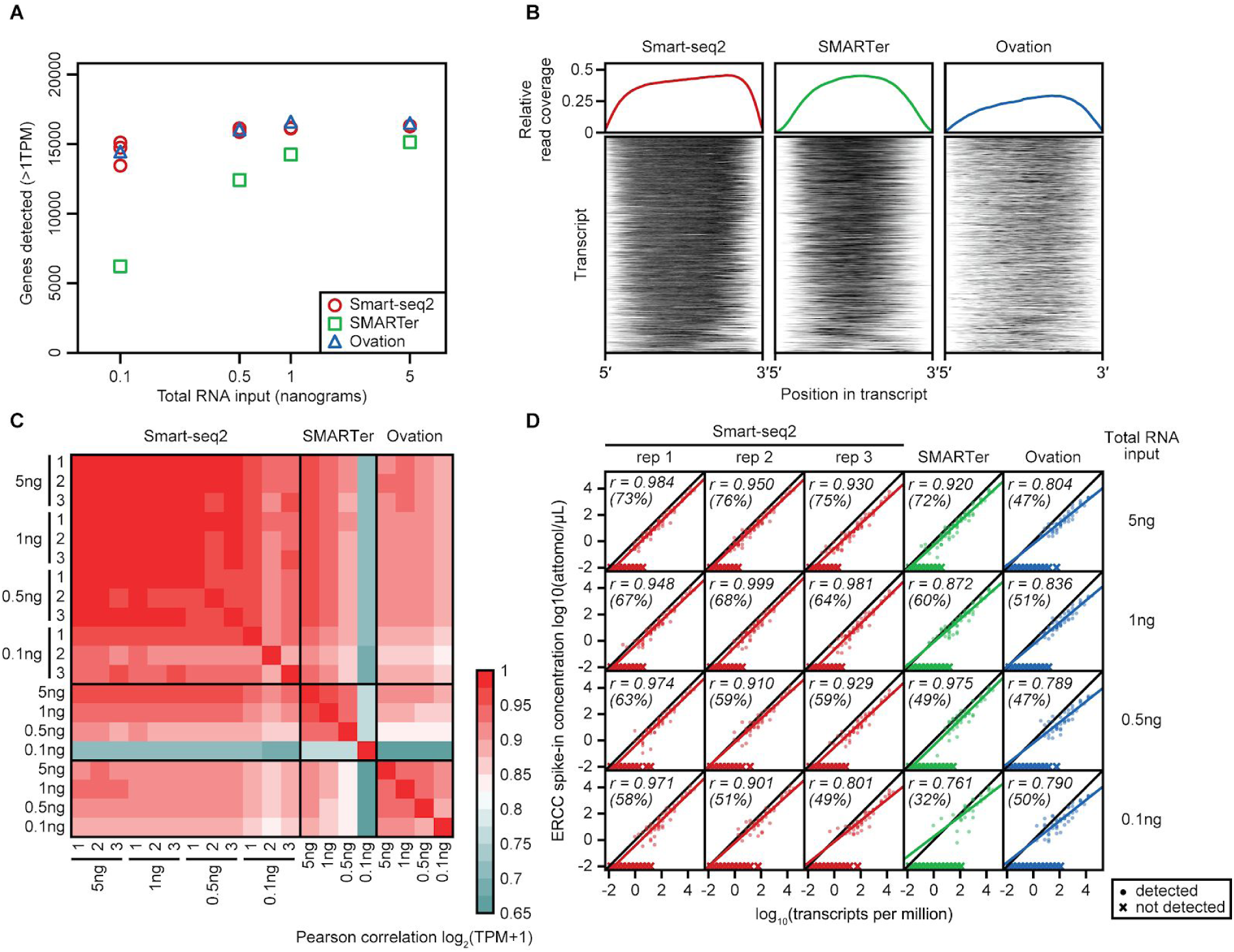
Comparison of different low-input mRNA-seq methods **(A)** Number of genes detected when using three different mRNA-seq library preparation methods from a dilution series of total RNA. **(B)** (Above) Read distribution along the length of all protein-coding transcripts for Smart-seq2, SMARTer, and Ovation samples generated from 100 pg of total RNA. Relative read coverage depth was binned into 100 bins from the 5′ terminus to 3′ terminus of each transcript. (Below) Heatmaps of read coverage for the 1000 most highly expressed transcripts across the three methods. **(C)** Heatmap depicting pairwise Pearson correlation of gene expression values for all samples in the dilution series. **(D)** Correlation of the TPM of all detected ERCC RNA spike-in molecules with their relative input concentration. r is Pearson correlation. Number in parentheses represents the percentage of ERCC spike-in oligos that were detectable in the given sample

We also assessed the reproducibility of quantifying transcript levels from varying amounts of low-input total RNA for the three mRNA-seq methods. Transcript levels across the dilution series were most highly correlated to each other for libraries generated with the Smart-seq2 protocol (Fig. 1C). The increased reproducibility of Smart-seq2 compared to the other two methods was most prevalent when starting with sub-nanogram levels of total RNA. To determine the accuracy and sensitivity of the three mRNA-seq library preparation methods, we introduced 92 synthetic poly(A) RNAs in specific molar ratios (i.e. ERCC spike-in mixes; LifeTech, (Baker et al. 2005)) to the samples before generating libraries with these methods. We compared the TPM levels detected for each ERCC spike-in with the relative amount added to each of the samples (Fig. 1D). Compared with the SMARTer and Ovation methods, libraries generated with Smart-seq2 consistently produced ERCC spike-in values that were more highly correlated with their known input amounts across the dilution series. Moreover, libraries generated with Smart-seq2 detected a higher percentage of the ERCC spike-ins added to the samples suggesting that Smart-seq2 was the most sensitive method. Altogether, our comparisons indicate that Smart-seq2 produces more sensitive, uniform, reproducible and accurate transcriptome profiles from Arabidopsis embryos especially when starting with sub-nanogram quantities of total RNA. Because the Smart-seq2 method is published, it also offers the advantage of being substantially less expensive compared with the other two methods.

### A developmental time series of Arabidopsis embryo transcriptomes

In order to profile transcriptome dynamics during Arabidopsis embryogenesis, we generated Smart-seq2 libraries from total RNA collected from embryos spanning presumptive morphogenesis (preglobular to late heart) and maturation (early torpedo to mature green) phases. More specifically, RNA was extracted from 50 embryos at each of these 8 different stages in biological triplicate (1,200 embryos total; Fig 2A). Embryos were staged based on their distinct morphologies as represented in Fig. 2A (see Materials & Methods for details regarding embryo isolations).

**Figure 2.**
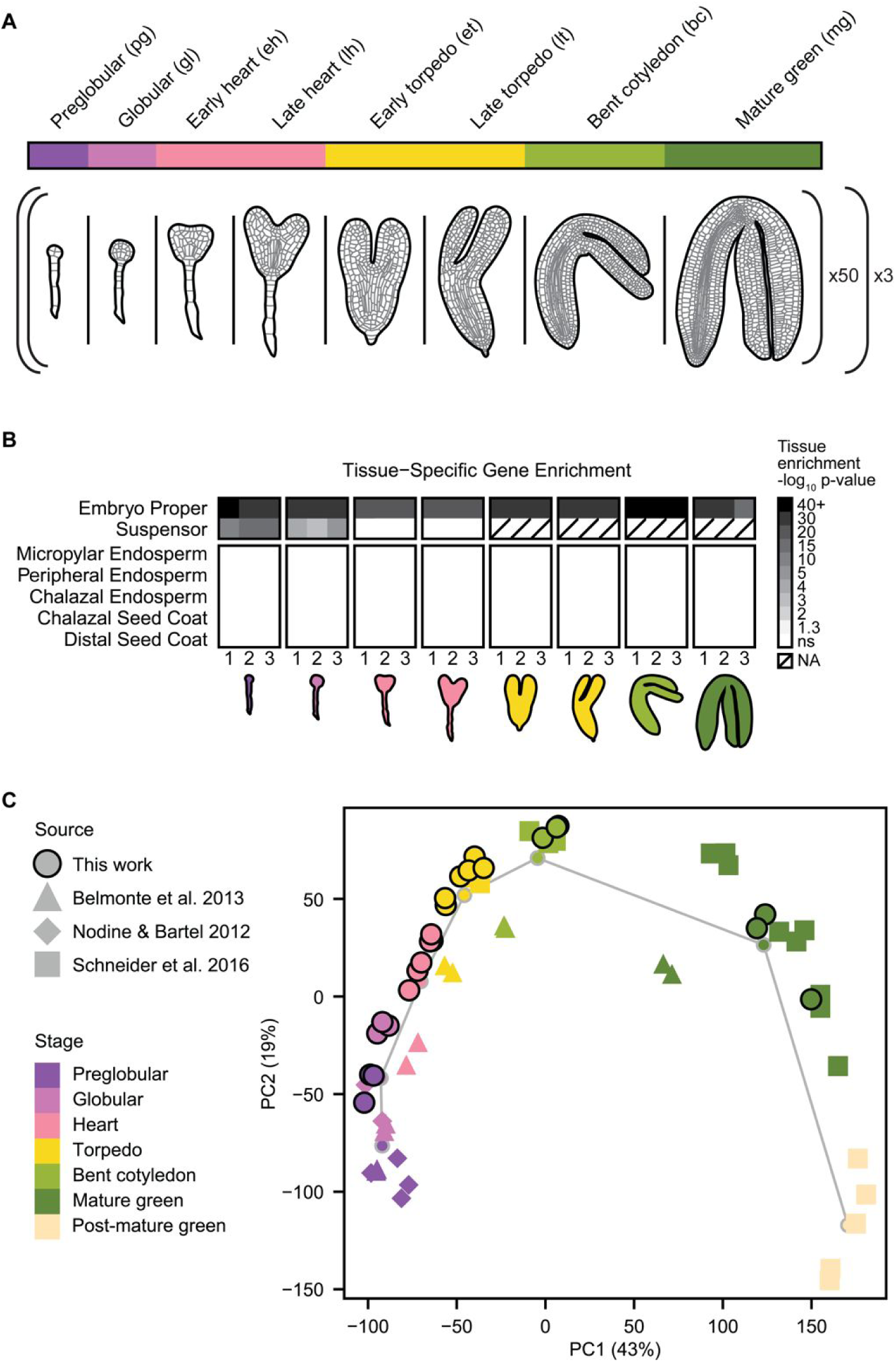
mRNA-seq time course of the Arabidopsis embryonic transcriptome **(A)** Overview of the performed experiment. From each of the displayed stages, total RNA was isolated from 50 embryos dissected from ovules in biological triplicate. Smart-seq2 libraries were prepared for each sample and the resulting libraries were sequenced on an Illumina instrument. **(B)** Results from the Tissue-Enrichment Test (Schon and Nodine 2017) on the obtained embryonic transcriptomes revealed a significant enrichment for embryo proper and suspensor transcripts (in the early stages), but no significant seed coat or endosperm contamination across all stages. **(C)** PCA displaying the embryo time series from this study in comparison with other embryo transcriptomics data (Nodine and Bartel 2012; Belmonte et al. 2013; Schneider et al. 2016). The centroids for each stage are depicted as dots connected by a gray line.

Sequencing of the 24 libraries on an Illumina sequencing platform yielded a total of over 792 million paired 50 base reads. After adaptor trimming with Cutadapt (Martin 2011), transcript abundances were quantified using Kallisto (Bray et al. 2016) and the Araport11 annotations (Cheng et al. 2017). In total, 624 million read pairs (78.8%) pseudoaligned concordantly to the Arabidopsis transcriptome (Online Table S1 & S2). Over 15,000 genes were detected at >1 TPM in every sample, with a gradual increase in total number of expressed genes through the early heart stage. We defined a gene as being expressed in the embryo if it has a mean TPM value of >1 in at least one developmental stage. Using this definition, we found that 21,433 genes (63.8% of all annotated genes) are expressed in developing embryos. Pearson correlations between biological replicates exceeded 0.97 in all pairwise comparisons, demonstrating that the results were highly reproducible (Online Table S3).

As previously shown, contamination by RNAs originating from the surrounding maternal seed coat and endosperm has been a frequent problem leading to erroneous conclusions when generating early embryonic transcriptomes (Schon and Nodine 2017). To determine whether significant RNA contamination exists in our datasets, we applied the Tissue-Enrichment Test (Schon and Nodine 2017) to each of the 24 embryonic sequencing libraries. This test revealed a strong enrichment of genes specific to the embryo proper in all samples (Fig. 2B). Additionally, suspensor-specific genes were statistically enriched in preglobular and globular embryos (adjusted p-value < 0.001). In contrast, none of the five seed coat or endosperm tissues were enriched in any sample at any stage. Therefore, we concluded that the mRNA-seq time series generated in this study represents transcripts exclusively from whole embryos.

To further assess the quality of our mRNA-seq time series, we compared it to publicly available embryonic gene expression data. The datasets we used were the laser capture microdissection (LCM) microarray data (Belmonte et al. 2013), an mRNA-seq time course of late stage embryos (Schneider et al. 2016) and timed, reciprocal crosses of early embryonic stages (Nodine and Bartel 2012). Each of these datasets were high quality and contamination-free (Online Fig. S1). To enable comparisons of the mRNA-seq and microarray datasets, we linearized the mRNA-seq data via DESeq2’s variance stabilizing transformation (VST) (Love et al. 2014) (Online Table S2). We then set a uniform baseline score of five for all datasets and performed a principal component analysis (PCA) (Fig. 2C). This analysis showed that PC 1 and 2, accounting together for about 64% of the variation, stratified the transcriptomes according to their developmental stage. The samples followed an archlike pattern, with transcriptomes from similar embryonic stages, but produced by different researchers, grouped in close proximity. This demonstrates that despite different methods of tissue isolation and transcript measurement, the global temporal dynamics revealed in the current study is consistent with expectations established by previously published datasets.

### Arabidopsis embryos have a unique transcriptome

To assess the broader developmental context of our embryo time series, we compared it to a large collection of transcriptomes from different Arabidopsis tissues (Klepikova et al. 2015, 2016). These two datasets currently provide the most comprehensive mRNA-seq maps available in Arabidopsis and include 27 different tissues and 31 developmental time points for a total of 153 distinct samples. We also included leaf and floral bud Smart-seq2 data that we previously generated to control for differences between protocols and lab conditions (Lutzmayer et al. 2017; Schon et al. 2018). Together with the early and late embryo samples described above, we have 186 additional mRNA-seq libraries to compare with the 24 embryo time series samples produced in this study.

To compare the embryonic transcriptome to post-embryonic tissues, we built a pairwise correlation matrix of the 210 samples mentioned above and included two RNA-seq collections generated on somatic embryos produced by two different protocols (Wickramasuriya and Dunwell 2015; Magnani et al. 2017); Online Table S3). This correlation matrix was used to hierarchically cluster all samples in a single dendrogram (Fig. 3A, all sample names shown in Online Table S1). Similar tissues largely grouped together, and a set of 15 general “tissue clusters” emerged, excluding samples derived from tissue culture. Embryo samples were partitioned into four clusters that were separated by developmental time: 1-cell/2-cell to globular stage embryos (referred to as pre-cotyledon), early heart through bent cotyledon (transition), green embryos 10 to 13 days after pollination (mature green), and post-green embryos 15 days after pollination or older clustered with dry seeds (post-mature green). These four groups will be referred to as embryonic “phases” below.

**Figure 3.**
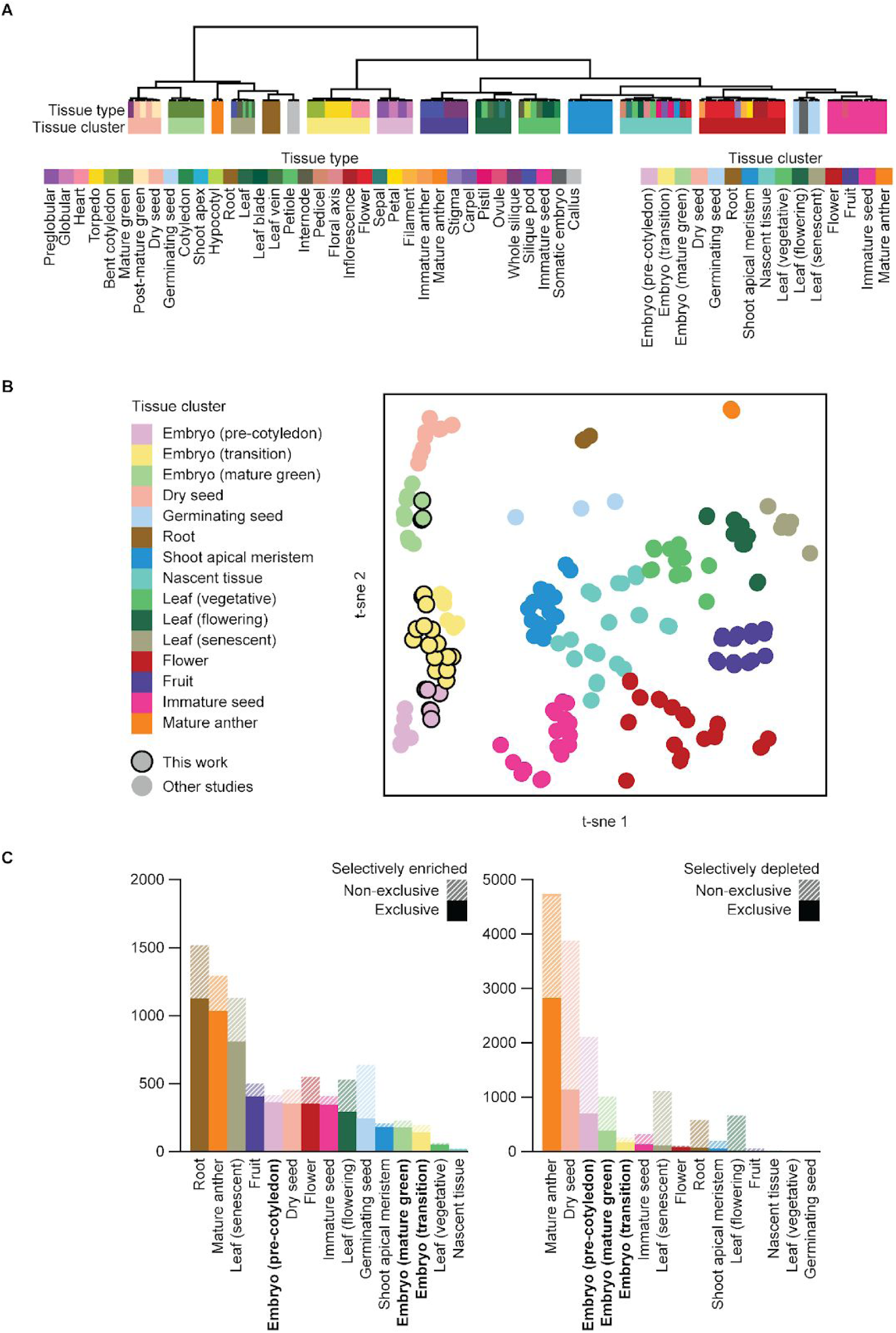
Comparison of embryo transcriptome to other plant tissues **(A)** Hierarchical clustering of 217 mRNA-seq libraries of various Arabidopsis tissues. Dendrogram was produced by clustering Pearson correlation of log_2_(TPM+1) using Ward’s criterion. **(B)** T-SNE comparing transcriptomes from different tissues and sources (Nodine and Bartel 2012; Belmonte et al. 2013; Klepikova et al. 2015, 2016; Schneider et al. 2016; Lutzmayer et al. 2017; Schon et al. 2018). Tissue clusters are defined by the results in panel **A**. **(C)** Number of genes in each tissue group cluster that show either enriched or depleted expression across different tissues. Minimum 4-fold change in tissue cluster vs. all other samples, ANOVA p-value <0.05 with Benjamini-Hochberg multiple testing correction. Gene expression was additionally required to be significantly higher than the neighboring set of samples on the hierarchical tree (ANOVA, BH-adjusted p-value <0.05)

To gain insights into the global relationships between these tissue clusters we performed t-distributed stochastic neighbor embedding (t-SNE) (Fig. 3B; detailed version Online Fig. S3). Post-embryonic tissue clusters radiated out from the “shoot apical meristem” cluster, connected via a loose collection of intermediate tissues that were labelled “nascent tissue”. In contrast, embryos formed a distinct group from all post-embryonic tissues and were stratified by developmental time. In both hierarchical clustering and t-SNE analysis, a large gap separates all embryos collected at or before eight days after pollination (bent cotyledon stage) from mature green embryos and dry seeds. Therefore, our analysis suggests that the Arabidopsis embryonic transcriptome undergoes radical global changes after both the globular and bent cotyledon stages.

To determine what specific transcript populations make the embryonic transcriptome unique, gene expression within all 15 tissue clusters was then compared to the global average. Transcripts that were at least 4-fold significantly more abundant (ANOVA, Benjamini-Hochberg adjusted p-value < 0.05) were considered enriched in that cluster if they were also significantly higher than their nearest neighbor on the dendrogram (Online Table S4). The inverse was also calculated to generate a list of genes depleted in a given tissue (Online Table S5). Roots had the highest number of genes enriched only in the given tissue and nowhere else (Fig 3C). Of embryo phases, pre-cotyledon possessed the largest number of exclusively enriched genes at 363. This set includes well-known embryo markers such as *LEC1, LEC2, PLT1, PLT2, WOX2, WOX8,* and *DRN* (Meinke et al. 1994; West et al. 1994; Haecker et al. 2004; Aida et al. 2004; Chandler et al. 2007; Galinha et al. 2007; Lau et al. 2012), and a much larger cohort of uncharacterized genes (Online Table S4 & S5). However, the most substantial difference between embryos and other plant tissue types appears to come from the genes specifically downregulated. With the exception of mature pollen, all four phases of embryogenesis had a larger set of genes exclusively depleted from their transcriptomes than any post-embryonic tissue. Additionally, 38% of genes depleted in the pre-cotyledon phase are also depleted in dry seeds. These genes are largely related to photosynthesis, including 13/13 annotated subunits of photosystem I, 9/15 subunits of photosystem II, and 14/20 light-harvesting complex genes. These core components of photosynthesis are absent early in morphogenesis, become abundant during the transition phase, but drop precipitously again in mature embryos (Online Fig. S3). This suggests that gene expression dynamics are tightly controlled during embryo development.

### Model based clustering reveals co-expressed groups, recapitulating specific biological functions

To examine temporal changes in gene expression, protein-coding genes detected at ≥1 TPM in at least one embryonic stage of the Smart-seq2 time series were subjected to model-based clustering by Mclust ((Scrucca et al. 2016), see Methods). Mclust classifies points according to a set of Gaussian Mixture Models and chooses the optimal model parameters to maximize the Bayesian Information Criterion (BIC). The mean TPM of 18,600 genes were converted to z-scores and based on analysis with Mclust, the optimal cluster number was 24 (Online Fig. 5). These 24 distinct covariance clusters, each composed of 290-1669 genes, were organized into four groups: (A) highest during the pre-cotyledon phase, (B) decreasing between transition and mature phases, (C) highest during the transition phase, and (D) highest during the mature phase (Fig. 4A; Online Table S6). To evaluate the accuracy of these covariance clusters, we examined the levels of 92 synthetic poly(A) RNAs (ERCC RNA Spike-In Mix; LifeTech, (Baker et al. 2005)) that were added to the samples in specific amounts during RNA isolation. The concentrations of the oligos in the mix spanned six orders of magnitude, but the ratio of each oligo relative to the others in the mix remained constant in all samples. Therefore, the ERCC spike-in RNAs represent a group of “covarying transcripts” across the time series. Fifty-seven ERCC RNAs passed the detection threshold of 1 TPM, covering four orders of magnitude of abundance. All 57 detected spike-in transcripts were placed in cluster A6, demonstrating that BIC clustering successfully identifies groups of covarying genes over a large dynamic range of expression.

**Figure 4.**
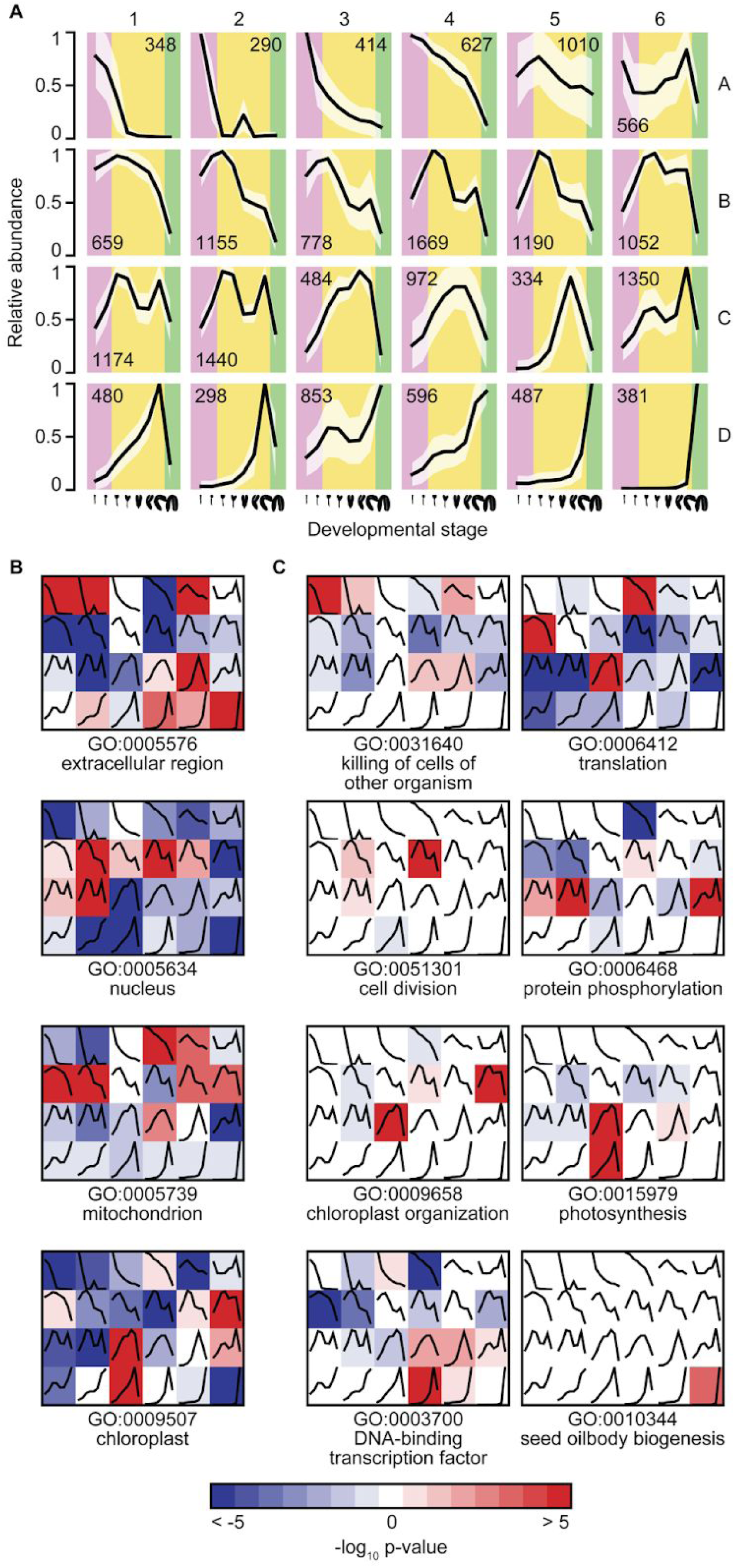
Covariance clustering of expressed genes across the embryo time series (**A)** Average maximum-normalized expression values for all genes in each of the 24 clusters generated by Mclust. Numbers in each panel indicate the number of genes grouped in that cluster, and polygons represents ±1 standard deviation. **(B)** Heatmaps of Gene Ontology (GO) term enrichment (red) and depletion (blue) for all 24 clusters for the terms “extracellular region”, “nucleus”, “mitochondrion”, and “chloroplast”. **(C)** Select GO term enrichments representative of the strongest enrichments across 24 clusters

**Figure 5.**
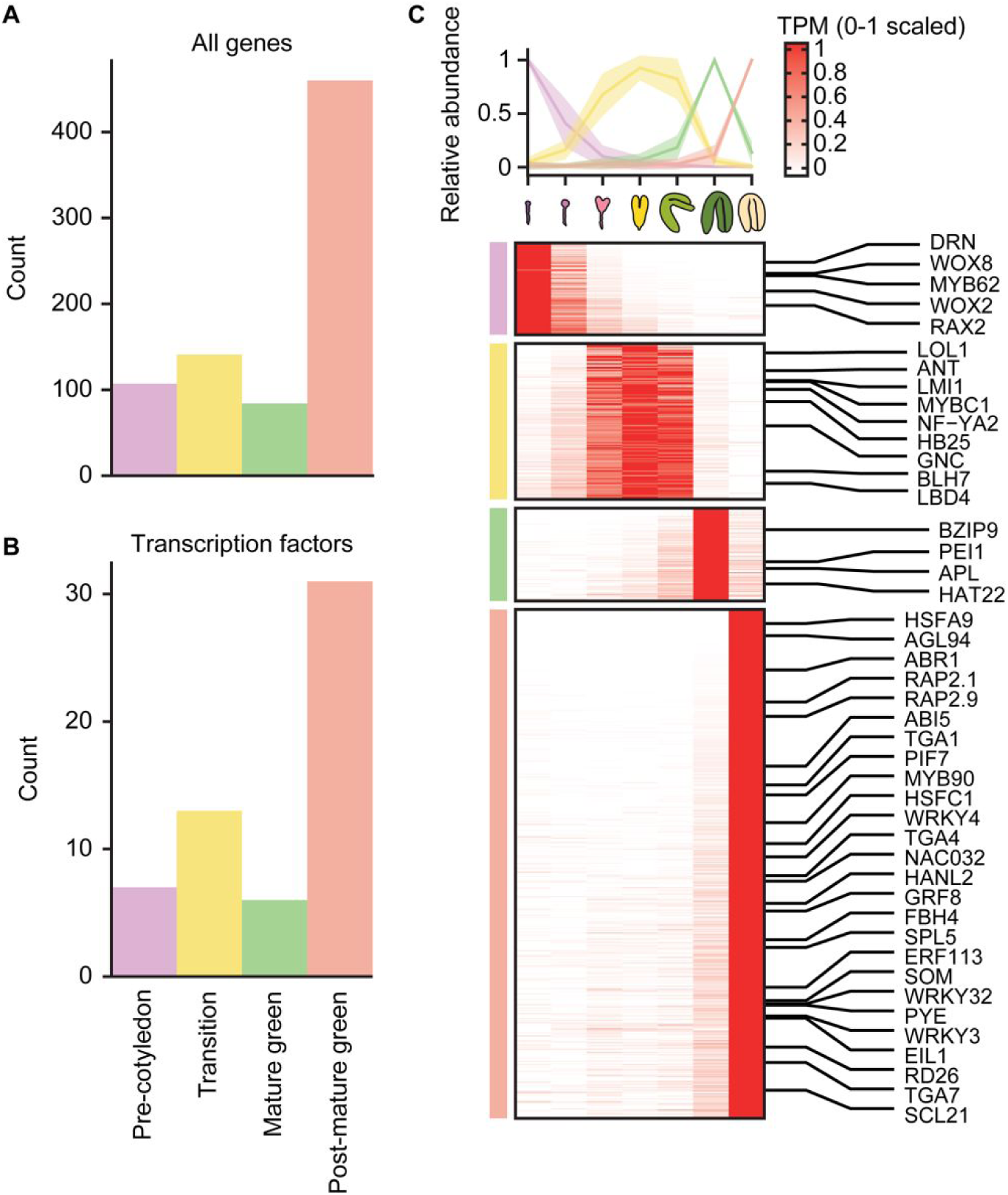
Identification of developmental phase markers Overview of the number of all **(A)** and transcription factor **(B)** marker genes identified for each developmental phase. **(C)** Marker gene transcript levels. (*Top*) Metaplots showing transcript levels for each marker group. Individual transcripts were normalized as TPM and scaled between 0 (not detected) and 1 (highest transcript levels observed). Purple, yellow, green and rose are used to indicate pre-cotyledon, transition, mature green or post-mature green phase embryos, respectively. Standard deviations for each set are indicated by the corresponding shading. (*Bottom*) A heatmap displaying row-scaled TPM values of marker genes identified (rows) across various embryonic stages (columns). Rows are sorted by their specificity score (decreasing). Phases are indicated by the colored bars on the left. Select transcription factors are labelled.

To investigate whether temporal covariance of gene expression in the embryo is connected to biological function, we performed Gene Ontology (GO) enrichment analysis on each BIC cluster (Ashburner et al. 2000; The Gene Ontology Consortium 2017). Compared to a background of all genes expressed during embryogenesis, all clusters showed an overrepresentation of at least one GO term (hypergeometric distribution p-value <0.05, Benjamini-Hochberg multiple testing correction; Online Table S6). A selection of the most strongly enriched terms in each cluster was examined in greater detail. For general insights on all clusters, enrichments and depletions for terms in the domain “cellular_component” are shown (Fig. 4B). Clusters peaking in the globular or heart stage have a tendency to be enriched for “mitochondrion” annotations, while “chloroplast” GO terms were predominantly associated with clusters that peak later in the transition toward mature embryos. In contrast, “nucleus” affiliated genes tend to be broadly expressed with a bias toward early expression, while genes with very early or very late expression tend to be associated with “extracellular region” GO terms.

The strongest early expression clusters were both enriched for the biological process term “killing of cells of other organism” (Fig. 4C). All 67 genes associated with this term in clusters A1 and A2 are defensin-like genes, a large family of short cysteine-rich peptides that were first characterized for their role as antimicrobial peptides (Silverstein et al. 2005), and a subset were later demonstrated to influence suspensor elongation during early embryogenesis (Costa et al. 2014). At the opposite end of the time series, cluster D6 is enriched in “seed oil body biogenesis” GO terms, represented by the oleosins *OLE1, OLE2,* and *OLE4*, as well as *SEIPIN1* and *OIL BODY-ASSOCIATED PROTEIN 1A*, both of which regulate the formation of the lipid droplets that accumulate in mature seeds (López-Ribera et al. 2014; Cai et al. 2015). Genes associated with “cell division” GO terms tend to be expressed throughout the time series but are highest at the early heart stage (cluster B4, Fig. 4C). This cluster is also enriched for embryo lethal mutations as defined by the Seedgenes database (Meinke et al. 2008), with 55 Seedgenes mutants belonging to this cluster (p-value < 2.3e^-8^, Online Fig. S6). This cluster is most strongly associated with the biological process “cell division” (GO:0051301, p-value < 2.2e^-14^) and contains 13 cyclins, along with 48 other cell cycle associated genes. Even more strongly enriched for embryo lethal genes is cluster B6, which peaks early during the heart stage but is sustained throughout the transition phase. This cluster is one of a few associated with “chloroplast organization”, and 28 of the 44 embryo lethal genes with this expression pattern are reported to arrest at the globular stage, including the gene *ACCUMULATION OF PHOTOSYSTEM ONE 2* (*APO2*, AT5G57930; (Meinke et al. 2008)). Indeed, a study focused on chloroplast-localized lethal genes concluded that embryo arrest at the globular stage was a common feature of chloroplast disruption (Bryant et al. 2011). Altogether this suggests that making photosynthetically active chloroplasts is a key checkpoint during embryo development, required to move beyond the morphogenesis phase.

### Identification of embryo specific marker transcripts

While forward genetic screens and functional studies have led to the identification of important developmental regulators and embryonic markers (Haecker et al. 2004; Meinke et al. 2008; Rademacher et al. 2011; Lau et al. 2012), we hypothesized that we could define additional markers with the embryonic transcriptome datasets. To identify embryonic markers we applied the tool MGFR (El Amrani et al. 2015) and treated different embryonic samples belonging to the four developmental phases (i.e. pre-cotyledon, transition, mature green and post-mature green). We required that protein-coding transcripts were >5 TPM in at least one embryonic stage, and after initial marker identification we also required that the marker transcript levels were at least five-fold higher compared to the other stages. This led to the identification of 107 pre-cotyledon, 141 transition, 84 mature green and 460 post-mature green marker transcripts (792 total; Fig. 5 A, B, Online Table S7). Because transcription factors are major determinants of cellular differentiation and have been utilized as embryonic markers, we investigated how many transcription factors are contained within this set of phase-enriched markers. For this we used the transcription factor annotations available from PlantTFDB 4.0 (Jin et al. 2017). We identified 7 pre-cotyledon, 13 transition, 6 mature green and 31 post-mature green marker transcription factors highly enriched in their respective phases (57 total; Fig. 5 A, B). In addition to identifying known transcription factor markers during morphogenesis such as *WOX2*, *WOX8* and *DRN*, we also identified new markers including two storekeeper protein-related transcripts (AT1G11510 and AT4G00390), AT1G68320/*MYB62* and AT2G36890/*RAX2* (Fig. 5C, Online Table S7). In addition to transcription factors, we also detected transcripts encoding a potential transcriptional co-activator (AT5G09240), a putative ubiquitin E3 ligase (AT3G11600) and an ubiquitin-like protein (AT1G53930). To provide a more complete resource, we also performed this marker analysis on the individual embryo stages (Online Table S7).

### Somatic and zygotic embryo transcriptomes are distinct from each other

Somatic embryos are widely thought to be suitable models for studying zygotic embryogenesis. However, it remains to be determined how similar somatic and zygotic embryos are to each other in terms of their transcript and protein populations. We hypothesized that transcripts up-regulated during the onset or progression of somatic embryogenesis should resemble zygotic embryo phases if somatic and zygotic embryo transcriptional processes are similar.

Two studies performed RNA-seq on somatic embryos derived from late-staged zygotic embryos during either their initiation (Magnani et al. 2017) or developmental progression (Wickramasuriya and Dunwell 2015). Magnani et al. collected torpedo staged embryos and dedifferentiated them into calli by culturing in auxin-rich medium in the dark. To induce somatic embryogenesis, callus cultures were then moved to auxin-free media and further cultured in dark conditions to induce somatic embryogenesis. *LEC2* expression is one of the earliest markers of calli that are competent to undergo somatic embryogenesis (Su et al. 2009). Thus, Magnani et al. purified nuclei from *LEC2* expressing calli cells (+LEC2) with INTACT (Deal and Henikoff 2010) to profile somatic embryo transcriptomes upon their initiation. RNA-seq was then performed on the +LEC2 samples together with isolated -LEC2 samples from the calli which served as a negative control. In the Wickramasuriya study, direct somatic embryogenesis was performed on bent cotyledon embryos whereby embryos were cultured with auxin under long day conditions and somatic embryos that could be morphologically distinguished from the surrounding calli were harvested 5, 10 and 15 days after auxin treatment. We quantified transcript levels from these studies as described above (see also Methods).

To assess whether transcriptomes of somatic embryos derived from late-stage zygotic embryos resemble those from early zygotic embryos, late zygotic embryos or another developmental phase, we first reanalyzed the published data to determine expressed protein-coding genes (>1 TPM) that were upregulated >4-fold (i.e. upregulated DEGs). The data from Magnani and colleagues was analyzed with DESeq2 (Love et al. 2014), allowing a FDR of 5%. We detected 236 upregulated DEGs (Online Table S8) of which 185 (78%) overlapped with those reported in this study. Because the data published by Wickramasuriya and Dunwell does not include biological replicates, we regarded all expressed protein-coding genes with transcripts increased at least four-fold during somatic embryo development as significantly upregulated (i.e. 112 upregulated DEGs; Online Table S8).

To determine which tissues the upregulated DEGs from somatic embryos are predominantly expressed in, we performed a tissue-enrichment test (TissueEnrich; (Jain and Tuteja 2018)). As a reference, we used the embryo time series described in this study together with the Klepikova expression data (Klepikova et al. 2015, 2016), and leaf and floral bud mRNA-seq datasets generated with Smart-seq2 by our group (Lutzmayer et al. 2017; Schon et al. 2018). To improve the ability to detect more tissue-specific gene expression patterns, the Klepikova data was manually curated to remove organs composed of tissues also sequenced at the same developmental stage (Online Table S8). Upon analyzing the tissue-enrichments of the Wickramasuriya upregulated DEGs, we observed a significant enrichment for nine non-embryonic tissues. Seven corresponded to shoot-derived tissues, and two were root tissues (Fig. 6 A). Similarly, the tissue-enrichment analysis of the Magnani DEGs also revealed no significant enrichment for embryonic tissues. However, the top two significantly enriched tissue types were root tissues (Fig. 6 B). We also repeated the analysis with the list of DEGs included in the original publications, and although we observed a trend towards a significant enrichment of heart stage embryos (p=0.06), no embryonic tissues passed the p<0.05 threshold (Online Fig. S6 & Online Table S8). In contrast, when the enrichment test was performed on upregulated DEGs from previously published early (Nodine and Bartel 2012) and late zygotic embryo datasets (Schneider et al. 2016), we detected a highly significant enrichment for the respective embryonic stages as expected (Online Fig. S6).

**Figure 6.**
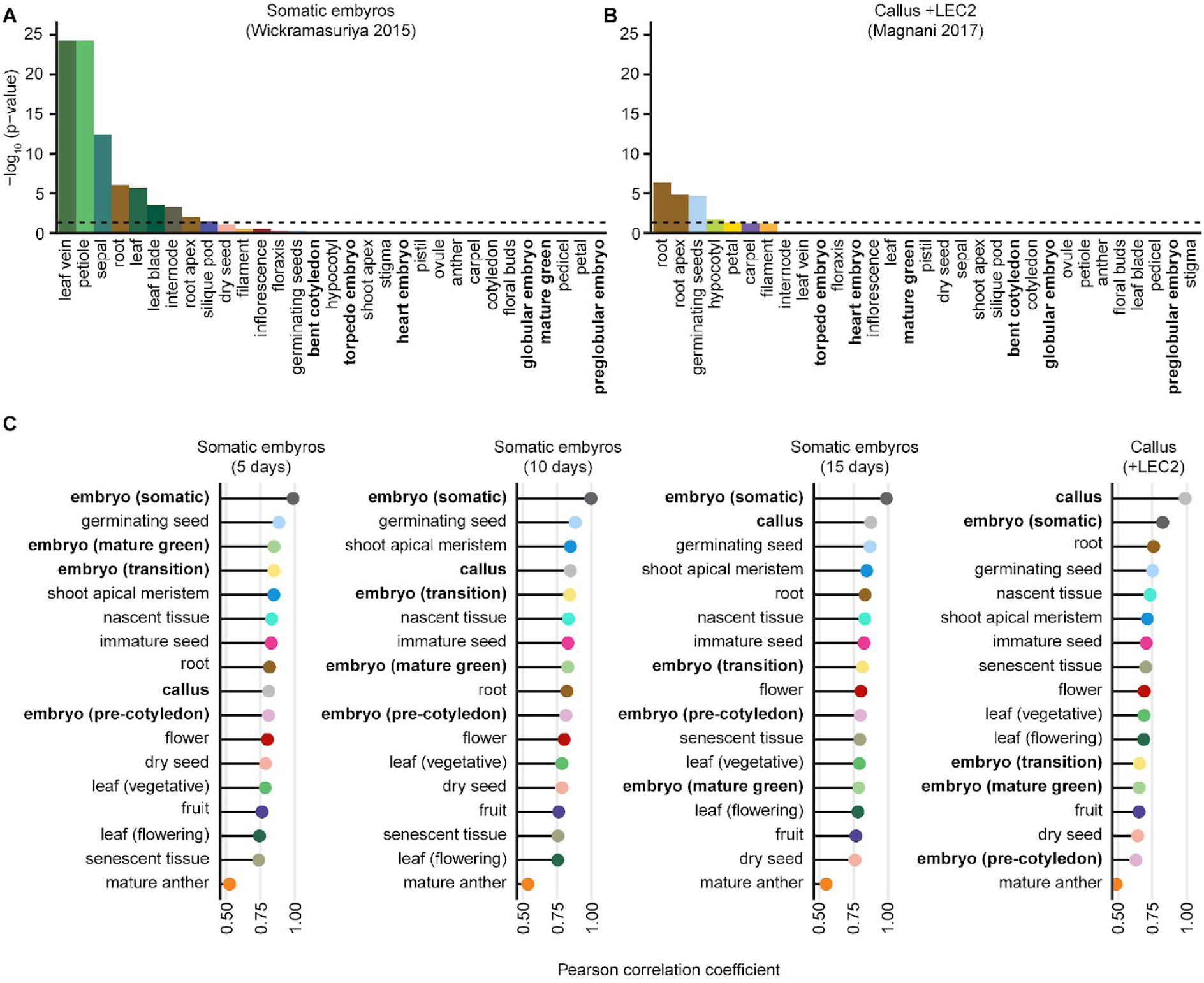
Somatic embryogenesis has limited resemblance to zygotic embryogenesis Tissue-specific gene enrichment (Jain and Tuteja 2018) of DEGs from somatic embryos (Wickramasuriya and Dunwell 2015) (**A**) and calli expressing *LEC2* (+LEC2) (Magnani et al. 2017) (**B**). DEGs were tested for enrichment against the embryo time series produced in this study and the Klepikova atlas (Klepikova et al. 2015, 2016). A significance level of p=0.05 is indicated by the dotted line. **(C)** Correlation analysis of the somatic embryo and LEC2+ transcriptomes against the tissue clusters established in Fig. 3. For a detailed overview which samples contribute to each cluster see **Online Table S1**.

In order to further estimate which tissues resemble somatic embryos derived from late-staged zygotic embryos, we performed a correlation analysis in which we compared their expression data against the mean expression of the previously identified tissue clusters (Fig. 6 C). Although somatic embryos are well-correlated with transcripts characteristic of transition and mature phases at their earliest time point (r=0.85), this similarity decreases over time. Moreover, as somatic embryos develop they appear to more resemble calli, as well as more specific tissues such as the shoot apical meristems and roots. We detected a very low correlation between the LEC2+ transcriptomes and all zygotic embryo stages. In fact, transcript abundances during the pre-cotyledon phase had the second lowest correlation of all 17 tissue clusters. When looking at a few select transcription factors (Lau et al. 2012) in more detail we also observed conflicting trends. We found that transcripts encoding several key developmental regulators such *WOX2, WOX8, LEC1* and *LEC2* were lowly abundant in the somatic embryos (< 5 TPM), or not at all in the LEC2+ cells (< 1 TPM), while others such as the PLETHORA family of transcription factors (*PLT1,PLT2,PLT3* and *BBM*) were highly abundant (> 30 TPM) (Online Fig. S7). Furthermore, we found a high correlation between both somatic embryo datasets and germinating seeds.

## Discussion

A variety of low-input mRNA sequencing (mRNA-seq) methods have been developed for tissue-specific and single-cell sequencing (Chen et al. 2018). Here we performed a side-by-side comparison of three low-input mRNA-seq protocols on Arabidopsis embryos and evaluated their performance. Our analysis showed that the substantially less-expensive Smart-seq2 method using off-the-shelf reagents significantly outperformed two commercially available kits when applied to low-input plant embryo RNA. We used Smart-seq2 to profile the transcriptomes of eight stages spanning embryonic development. Our data are consistent with other published transcriptomes and bridges an important gap previously missing in the field. While other studies were able to profile either early or late Arabidopsis embryos, we obtained a more comprehensive time series, from the preglobular to the mature green stages. Our analysis has shown that these transcriptomes are of high quality and free of contamination from maternal tissues. Moreover, because the embryonic transcriptomes presented here were generated with Smart-seq2 technology and deeply sequenced, they also have an increased number of detectable genes with more uniform coverage along the transcripts, and a larger dynamic range relative to other early embryonic datasets.

We observed that embryos have a unique transcriptome compared to other Arabidopsis tissue types. We speculate that this is due to the unique differentiation processes occurring during embryogenesis. This is supported by the results of our model-based clustering analysis, which indicates that different biological processes are enriched during the four different phases of embryogenesis. For example, model-based clustering correctly i) co-clustered all ERCC spike-in controls, ii) identified a functional enrichment of mitochondria related transcripts in the pre-cotyledon stages (Gao et al. 2018), and iii) also correctly detected timing of accumulation of chlorophyll accumulation (Kim et al. 2002).

Analysis of the embryonic transcriptomes produced in this and other studies indicates that Arabidopsis embryonic development can be partitioned into either pre-cotyledon, transition, mature green or post-mature green phases, each of which are characterized by distinct biological processes. Based on these results, we propose that embryos progress through these four distinct phases of development prior to the onset of germination. We were also able to establish a set of stringently defined temporal markers. In addition to several known important developmental regulators, this set of stage-specific markers contains many uncharacterized candidates for follow-up gene expression and mutagenesis studies.

Although somatic embryos are often thought to be a suitable model to study gene-regulatory processes occurring during zygotic embryogenesis, we observed significant differences between the transcriptomes of zygotic and somatic embryos. Based on our analysis, this appears to be at least partially due to the culturing conditions used to generate somatic embryos. For example, we observed a significant enrichment for green tissues in the DEG set from the Wickramasuriya et al. study (long day light conditions), and a predominant enrichment of non-green tissues in the DEGs from Magnani et al. (cultured in dark). Furthermore, we detected a strong correlation of both somatic embryos and LEC+ cells with germinating seeds, which tentatively suggest that somatic embryogenesis may more closely resemble processes occurring during germination rather than embryogenesis. However, we could not detect the expression of several select transcription factors in the somatic embryo datasets including *LEC2* transcripts which were <1 TPM in the LEC2+ cells. Therefore, our analysis suggests that zygotic and somatic embryos are transcriptionally distinct. However, the field would benefit from further transcriptome comparisons between zygotic embryos and additional datasets of somatic embryos derived from either late-staged zygotic embryos or explants from mutants or stress-treated tissues (Mozgová et al. 2017; Kadokura et al. 2018).

## Materials & Methods

### Plant material and growth

Col-0 seeds were grown in a climate controlled growth chambers set at 20-22° C temperature with a 16h light/8h dark cycle.

### RNA extraction, cDNA library preparation and next-generation sequencing

Embryos were dissected as described in (Nodine and Bartel 2010) except that embryos were dissected and washed 3× in 10% RNAlater (ThermoFisher). RNA was isolated from 50 embryos per sample collected in approximately 30 µl of 100% RNAlater by adding 500 µl of TRIzol (Life Tech) followed by brief vortexing and incubating at 60° C for 30 minutes. Sterile nuclease-free pestles were used to crush bent-cotyledon and mature green staged embryos (50×) within a 1.5 ml tube. ERCC spike-ins (LifeTech) were added during the TRIzol preparation after the addition of chloroform. Precipitated RNA was resuspended with 5-12 µl of nuclease-free water, and 1 µl was used for mRNA-seq library construction. mRNA-seq libraries were prepared with SMARTer Ultra Low Input RNA Kit for sequencing - v3 (Clontech) or Ovation PicoSL WTA System V2 (Nugen) according to the manufacturer’s recommendations. Smart-seq2 libraries were generated according to (Picelli et al. 2013). To control for library quality, length distributions of both amplified cDNA and final libraries were inspected using an Agilent DNA HS Bioanalyzer Chip. Libraries were diluted and sequenced with paired-end 50 base mode on an Illumina HiSeq 2500 machine.

### Pseudo-alignment & mRNA-seq quantification

The pseudoaligner Kallisto was used for quantification of all mRNA-seq datasets (v0.44.0, (Bray et al. 2016)). An index was generated for all transcripts in the Ensembl build of the TAIR10 annotation set (release version 40, ftp://ftp.ensemblgenomes.org/pub/plants/release-40/gff3/arabidopsis_thaliana/Arabidopsis_thaliana.TAIR10.40.gff3.gz), including all 92 ERCC RNA spike-in sequences (https://www-s.nist.gov/srmors/certificates/documents/SRM2374_putative_T7_products_NoPolyA_v1.fasta). First, a FASTA file containing each transcript model was built by running bedtools getfasta (v2.17.0, (Quinlan and Hall 2010)) with the TAIR10 GFF3 above and the TAIR10 genome (ftp://ftp.ensemblgenomes.org/pub/plants/release-40/fasta/arabidopsis_thaliana/dna/Arabidopsis_thaliana.TAIR10.dna.toplevel.fa.gz). Then, *kallisto index* was run on the transcript FASTA file to generate an index file.

Before quantification, the appropriate adapter sequences for each mRNA-seq library were trimmed from the FASTQ files using Cutadapt with a minimum match length of five nucleotides (v1.9.1, (Martin 2011)). Cutadapt was also used to trim all oligo-A or oligo-T sequences that were at least five nucleotides long from the ends of reads. All reads longer than 18 nucleotides after trimming were used as input for *kallisto quant*. *kallisto quant* was run on paired-end samples using default settings. For single-end samples, the arguments *--fragment-length 200 --sd 100* were used. Gene-level transcripts per million (TPM) were estimated by combining the TPM of all isoforms of protein-coding genes. Gene IDs mapping to mitochondria and chloroplast genomes, as well as the 270 kilobase mitochondrial insertion on chromosome 2 (Stupar et al. 2001), were discarded. Last, abundances were renormalized to a sum of 1 million for each sample.

### Quality control & tissue enrichment testing

Seed tissue enrichment tests were performed with the previously published *tissue-enrichment-test* (Schon and Nodine 2017) using default parameters and gene-level TPM tables described above as input. For comparison of mRNA-seq data to microarray data from (Belmonte et al. 2013), a table of raw mRNA-seq read counts mapping to each gene was combined with the mean-centered signal intensity scores from the Series Matrix File for GEO series GSE11262. Genes not represented on the Ath1 array by a single unambiguous prober were discarded. The samples in this table were normalized with the varianceStabilizingTransformation function of the *R* library *DEseq2* (Love et al. 2014). Values from this table were reduced by five, and all negative values were set to zero in order to set a uniform baseline between samples. Principal Component Analysis (PCA) was performed with the *R* function *prcomp(center = T, scale = F)*.

To identify genes enriched or depleted in a cluster of tissue samples, a hierarchical tree of all samples was first established. Pearson correlation of log_2_(TPM+1) was calculated between all pairs of samples, and hierarchical clustering was performed on this correlation matrix with the R library pheatmap, using *clustering_method = ‘ward.D’*. An ANOVA model was built with the *R* function *aov()* to compare each tissue cluster to its nearest neighbor and to the outgroup of all other samples. Any gene whose expression is significantly higher than the neighboring cluster (ANOVA p-value <0.05, Benjamini-Hochberg multiple testing correction), as well as significantly higher than the global expression with a minimum fold change of 4 and ANOVA adjusted p-value <0.05, is considered enriched for that tissue. In contrast, a gene is considered depleted for that tissue if it is significantly lower in expression than both the global average and the neighboring cluster is considered depleted for that tissue. If a gene is enriched in one tissue and no other tissues, it is considered “exclusively enriched” in that tissue. Likewise, exclusively depleted tissues are not significantly depleted in any other tissue cluster.

### Identification of marker genes

For the identification of phase-specific markers, we imported the TPM values from this study and previously published data (Nodine and Bartel 2012; Schneider et al. 2016) into *R* (v3.5.1). We then applied the *MGFR* (v1.6) tool (El Amrani et al. 2015) with default settings using these TPM values and treating different samples from the same developmental phase (pre-cotyledon, transition, mature green, post-mature green) as independent replicates. The resulting gene list was then subsetted for markers with a score lower than 0.2, which corresponds to a 5-fold increase in marker expression compared to its background (the respective other stages). The heatmap in Fig. 5 C was generated with the *ComplexHeatmap* package (Gu et al. 2016).

### Model-based clustering

The *R* library *Mclust* (Scrucca et al. 2016) was used to partition expressed genes into covariance clusters. First, a mean TPM value was calculated across the three biological replicates of each stage in the Smart-seq2 embryo time series. These mean TPM values were converted to z-score, or (*x_i_* - *x̄*)/*s*, where x_i_ is the TPM value for a gene in a stage, *x̄* is the mean of TPM values across all stages in the time series, and *s* is the standard deviation of TPM across all stages. The z-scores for all genes with a mean TPM of at least 1 in ≥1 stage were analyzed with the function *mclustBIC(modelNames = “VVV”, G = seq(2,50,by=2))* to calculate the Bayesian Information Content (BIC) for models with 2 to 50 components.(Online Fig. S4). The first step-wise increase in the number of components that decreased BIC was chosen as the optimal number of components (24). Then the function *Mclust* was run with the settings *data = 24, modelNames = “VVV”, prior = priorControl()*.

### Somatic embryo differentially expressed gene testing

First, Kallisto output files from either the Wickramasuriya and Dunwell, or the Magnani et al. study were imported into *R* (v3.5.1) with the *tximport* package (v1.8) (Soneson et al. 2015). The count data were then subsetted for nuclear protein-coding genes (see genes column in Online Table S2) and variance stabilized via *DESeq2* (v1.20) (Love et al. 2014). Identification of differentially expressed genes in the Wickramasuriya and Dunwell study was performed similar as in the original publication (Wickramasuriya and Dunwell 2015). We calculated the VST fold changes between the 5 and 10 day samples, and the 10 and 15 day samples, respectively. All genes with transcripts > 1 TPM at one stage and VST fold changes > 2 were kept as DEGs. For the Magnani et al. data we used *DESeq2’s* (v1.20) pairwise Wald-test (Love et al. 2014) to detect DEGs between the callus and the LEC2 intact purified callus nuclei (Magnani et al. 2017). DEGs where then further subsetted as described above (transcripts > 1 TPM, VST fold change > 2).

### Somatic embryo tissue enrichment testing

The TPM expression values of all samples were imported to *R* (v3.5.1) and then subsetted for our embryo time series and the Klepikova expression data. To avoid obfuscation of more specific gene expression patterns, the Klepikova data was manually curated to remove composite tissues, that had more specific subtissues sequenced at the same developmental age (Online Table S8). We then used this data to train the *R* package *TissueEnrich* (v1.0.6) (Jain and Tuteja 2018) with default parameters. We then used this package to test in which tissues DEGs are predominantly enriched/overexpressed.

## Supporting information

## Author Contributions

M.D.N. performed the experiments, F.H. and M.A.S. analyzed the data, and all authors interpreted the results and wrote the manuscript.

## Conflict of Interest

The authors declare that they have no conflict of interest.

### Acknowledgements

We thank the VBCF NGS Unit for sequencing Smart-seq2 libraries, the VBCF Plant Sciences Facility for plant growth chamber access and the GMI High Performance Computing Group for computational resources and technical support. We furthermore thank Ulf Naumann for technical assistance with comparing mRNA-seq methods, as well as Ping Kao for his input on somatic embryo comparisons. We also thank Alexandra Plotnikova, Stefan Lutzmayer, Ruben Gutzat, Mattia Doná and Michael Borg for critical feedback on the figures.

## References

Aida M, Beis D, Heidstra R, et al (2004) The PLETHORA genes mediate patterning of the Arabidopsis root stem cell niche. Cell 119:109–120. doi: 10.1016/j.cell.2004.09.018

Ashburner M, Ball CA, Blake JA, et al (2000) Gene ontology: tool for the unification of biology. The Gene Ontology Consortium. Nat Genet 25:25–29. doi: 10.1038/75556

Baker SC, Bauer SR, Beyer RP, et al (2005) The External RNA Controls Consortium: a progress report. Nat Methods 2:731–734. doi: 10.1038/nmeth1005-731

Belmonte MF, Kirkbride RC, Stone SL, et al (2013) Comprehensive developmental profiles of gene activity in regions and subregions of the Arabidopsis seed. Proc Natl Acad Sci U S A 110:E435–44. doi: 10.1073/pnas.1222061110

Bray NL, Pimentel H, Melsted P, Pachter L (2016) Near-optimal probabilistic RNA-seq quantification. Nat Biotechnol 34:525–527. doi: 10.1038/nbt.3519

Bryant N, Lloyd J, Sweeney C, et al (2011) Identification of nuclear genes encoding chloroplast-localized proteins required for embryo development in Arabidopsis. Plant Physiol 155:1678–1689. doi: 10.1104/pp.110.168120

Cai Y, Goodman JM, Pyc M, et al (2015) Arabidopsis SEIPIN Proteins Modulate Triacylglycerol Accumulation and Influence Lipid Droplet Proliferation. Plant Cell 27:2616–2636. doi: 10.1105/tpc.15.00588

Chandler JW, Cole M, Flier A, et al (2007) The AP2 transcription factors DORNROSCHEN and DORNROSCHEN-LIKE redundantly control Arabidopsis embryo patterning via interaction with PHAVOLUTA. Development 134:1653–1662. doi: 10.1242/dev.001016

Cheng C-Y, Krishnakumar V, Chan AP, et al (2017) Araport11: a complete reannotation of the Arabidopsis thaliana reference genome. Plant J 89:789–804. doi: 10.1111/tpj.13415

Chen X, Teichmann SA, Meyer KB (2018) From Tissues to Cell Types and Back: Single-Cell Gene Expression Analysis of Tissue Architecture. Annu Rev Biomed Data Sci 1:29–51. doi: 10.1146/annurev-biodatasci-080917-013452

Costa LM, Marshall E, Tesfaye M, et al (2014) Central cell-derived peptides regulate early embryo patterning in flowering plants. Science 344:168–172. doi: 10.1126/science.1243005

Deal RB, Henikoff S (2010) A simple method for gene expression and chromatin profiling of individual cell types within a tissue. Dev Cell 18:1030–1040. doi: 10.1016/j.devcel.2010.05.013

El Amrani K, Stachelscheid H, Lekschas F, et al (2015) MGFM: a novel tool for detection of tissue and cell specific marker genes from microarray gene expression data. BMC Genomics 16:645. doi: 10.1186/s12864-015-1785-9

Galinha C, Hofhuis H, Luijten M, et al (2007) PLETHORA proteins as dose-dependent master regulators of Arabidopsis root development. Nature 449:1053–1057. doi: 10.1038/nature06206

Gao L, Guo X, Liu X-Q, et al (2018) Changes in mitochondrial DNA levels during early embryogenesis in Torenia fournieri and Arabidopsis thaliana. Plant J. doi: 10.1111/tpj.13987

Gu Z, Eils R, Schlesner M (2016) Complex heatmaps reveal patterns and correlations in multidimensional genomic data. Bioinformatics 32:2847–2849. doi: 10.1093/bioinformatics/btw313

Haecker A, Gross-Hardt R, Geiges B, et al (2004) Expression dynamics of WOX genes mark cell fate decisions during early embryonic patterning in Arabidopsis thaliana. Development 131:657–668. doi: 10.1242/dev.00963

Jain A, Tuteja G (2018) TissueEnrich: Tissue-specific gene enrichment analysis. Bioinformatics. doi: 10.1093/bioinformatics/bty890

Jin J, Tian F, Yang D-C, et al (2017) PlantTFDB 4.0: toward a central hub for transcription factors and regulatory interactions in plants. Nucleic Acids Res 45:D1040–D1045. doi: 10.1093/nar/gkw982

Kadokura S, Sugimoto K, Tarr P, et al (2018) Characterization of somatic embryogenesis initiated from the Arabidopsis shoot apex. Dev Biol 442:13–27. doi: 10.1016/j.ydbio.2018.04.023

Kim I, Hempel FD, Sha K, et al (2002) Identification of a developmental transition in plasmodesmatal function during embryogenesis in Arabidopsis thaliana. Development 129:1261–1272

Klepikova AV, Kasianov AS, Gerasimov ES, et al (2016) A high resolution map of the Arabidopsis thaliana developmental transcriptome based on RNA-seq profiling. Plant J 88:1058–1070. doi: 10.1111/tpj.13312

Klepikova AV, Logacheva MD, Dmitriev SE, Penin AA (2015) RNA-seq analysis of an apical meristem time series reveals a critical point in Arabidopsis thaliana flower initiation. BMC Genomics 16:466. doi: 10.1186/s12864-015-1688-9

Lau S, Slane D, Herud O, et al (2012) Early embryogenesis in flowering plants: setting up the basic body pattern. Annu Rev Plant Biol 63:483–506. doi: 10.1146/annurev-arplant-042811-105507

López-Ribera I, La Paz JL, Repiso C, et al (2014) The evolutionary conserved oil body associated protein OBAP1 participates in the regulation of oil body size. Plant Physiol 164:1237–1249. doi: 10.1104/pp.113.233221

Love MI, Huber W, Anders S (2014) Moderated estimation of fold change and dispersion for RNA-seq data with DESeq2. Genome Biol 15:550. doi: 10.1186/s13059-014-0550-8

Lutzmayer S, Enugutti B, Nodine MD (2017) Novel small RNA spike-in oligonucleotides enable absolute normalization of small RNA-Seq data. Sci Rep 7:5913. doi: 10.1038/s41598-017-06174-3

Magnani E, Jiménez-Gómez JM, Soubigou-Taconnat L, et al (2017) Profiling the onset of somatic embryogenesis in Arabidopsis. BMC Genomics 18:998. doi: 10.1186/s12864-017-4391-1

Martin M (2011) Cutadapt removes adapter sequences from high-throughput sequencing reads. EMBnet.journal 17:10–12. doi: 10.14806/ej.17.1.200

Meinke D, Muralla R, Sweeney C, Dickerman A (2008) Identifying essential genes in Arabidopsis thaliana. Trends Plant Sci 13:483–491. doi: 10.1016/j.tplants.2008.06.003

Meinke DW, Franzmann LH, Nickle TC, Yeung EC (1994) Leafy Cotyledon Mutants of Arabidopsis. Plant Cell 6:1049–1064. doi: 10.1105/tpc.6.8.1049

Mozgová I, Muñoz-Viana R, Hennig L (2017) PRC2 Represses Hormone-Induced Somatic Embryogenesis in Vegetative Tissue of Arabidopsis thaliana. PLoS Genet 13:e1006562. doi: 10.1371/journal.pgen.1006562

Nodine MD, Bartel DP (2012) Maternal and paternal genomes contribute equally to the transcriptome of early plant embryos. Nature 482:94–97. doi: 10.1038/nature10756

Nodine MD, Bartel DP (2010) MicroRNAs prevent precocious gene expression and enable pattern formation during plant embryogenesis. Genes Dev 24:2678–2692. doi: 10.1101/gad.1986710

Picelli S, Björklund ÅK, Faridani OR, et al (2013) Smart-seq2 for sensitive full-length transcriptome profiling in single cells. Nat Methods 10:1096–1098. doi: 10.1038/nmeth.2639

Quinlan AR, Hall IM (2010) BEDTools: a flexible suite of utilities for comparing genomic features. Bioinformatics 26:841–842. doi: 10.1093/bioinformatics/btq033

Rademacher EH, Möller B, Lokerse AS, et al (2011) A cellular expression map of the Arabidopsis AUXIN RESPONSE FACTOR gene family: A cellular expression map of ARF gene expression. Plant J 68:597–606. doi: 10.1111/j.1365-313X.2011.04710.x

Schneider A, Aghamirzaie D, Elmarakeby H, et al (2016) Potential targets of VIVIPAROUS1/ABI3-LIKE1 (VAL1) repression in developing Arabidopsis thaliana embryos. Plant J 85:305–319. doi: 10.1111/tpj.13106

Schon MA, Kellner MJ, Plotnikova A, et al (2018) NanoPARE: parallel analysis of RNA 5’ ends from low-input RNA. Genome Res. doi: 10.1101/gr.239202.118

Schon MA, Nodine MD (2017) Widespread Contamination of Arabidopsis Embryo and Endosperm Transcriptome Data Sets. Plant Cell 29:608–617. doi: 10.1105/tpc.16.00845

Scrucca L, Fop M, Murphy TB, Raftery AE (2016) mclust 5: Clustering, Classification and Density Estimation Using Gaussian Finite Mixture Models. R J 8:289–317

Silverstein KAT, Graham MA, Paape TD, VandenBosch KA (2005) Genome organization of more than 300 defensin-like genes in Arabidopsis. Plant Physiol 138:600–610. doi: 10.1104/pp.105.060079

Soneson C, Love MI, Robinson MD (2015) Differential analyses for RNA-seq: transcript-level estimates improve gene-level inferences. F1000Res 4:1521. doi: 10.12688/f1000research.7563.2

Stupar RM, Lilly JW, Town CD, et al (2001) Complex mtDNA constitutes an approximate 620-kb insertion on Arabidopsis thaliana chromosome 2: implication of potential sequencing errors caused by large-unit repeats. Proc Natl Acad Sci U S A 98:5099–5103. doi: 10.1073/pnas.091110398

Su YH, Zhao XY, Liu YB, et al (2009) Auxin-induced WUS expression is essential for embryonic stem cell renewal during somatic embryogenesis in Arabidopsis. Plant J 59:448–460. doi: 10.1111/j.1365-313X.2009.03880.x

The Gene Ontology Consortium (2017) Expansion of the Gene Ontology knowledgebase and resources. Nucleic Acids Res 45:D331–D338. doi: 10.1093/nar/gkw1108

West M, Yee KM, Danao J, et al (1994) LEAFY COTYLEDON1 Is an Essential Regulator of Late Embryogenesis and Cotyledon Identity in Arabidopsis. Plant Cell 6:1731–1745. doi: 10.1105/tpc.6.12.1731

Wickramasuriya AM, Dunwell JM (2015) Global scale transcriptome analysis of Arabidopsis embryogenesis in vitro. BMC Genomics 16:301. doi: 10.1186/s12864-015-1504-6

