## Supplementary material for "The embryonic transcriptome of *Arabidopsis thaliana*"

**Plant Reproduction online resources for "The embryonic transcriptome of *Arabidopsis thaliana*"**

### Table of Contents

| | Page |
|---|---|
| **Online Methods** | **2** |
| DEG testing for additional early and late embryo samples | 2 |
| **Online Figures** | **3** |
| Figure S1 Tissue-enrichment-test for referenced embryo transcriptomes | 3 |
| Figure S2 t-SNE of all samples | 4 |
| Figure S3 Expression of photosynthesis pathway components during embryogenesis | 5 |
| Figure S4 BIC score for model based clustering | 6 |
| Figure S5 Gene group enrichment for model based clustering | 7 |
| Figure S6 Additional TissueEnrich results | 8 |
| Figure S7 Expression of select embryonic markers genes | 9 |
| **Online Tables** | **11** |
| Table S1 Sample overview | 11 |
| Table S2 Expression values | 11 |
| Table S3 Pearson correlation matrix | 11 |
| Table S4 Tissue-cluster enriched genes | 11 |
| Table S5 Tissue-cluster depleted genes | 11 |
| Table S6 Model based clustering | 11 |
| Table S7 Phase and stage specific markers | 11 |
| Table S8 DEGs and TissueEnrich results | 11 |
| **Online References** | **12** |

**Online Methods**

**DEG testing for additional early and late embryo samples**

For the identification of differentially expressed genes in the early (Nodine and Bartel 2012) and late embryos (Schneider et al. 2016) study *Kallisto* output files were read via the *tximport* package (v1.8) (Soneson et al. 2015) into R 3.5.1. The count data was then subsetted for nuclear, protein coding genes (see genes column in Online Table S2). We then applied *DESeq2's* (v1.20) (Love et al. 2014) likelihood-ratio test (*design = ~ time.point, reduced = 1*) independently to each dataset, allowing an FDR up to 5%. The resulting list of DEGs was subsetted for expressed (TPM > 1), upregulated genes ($\log_2$ fold-change >2).

### Online Figures

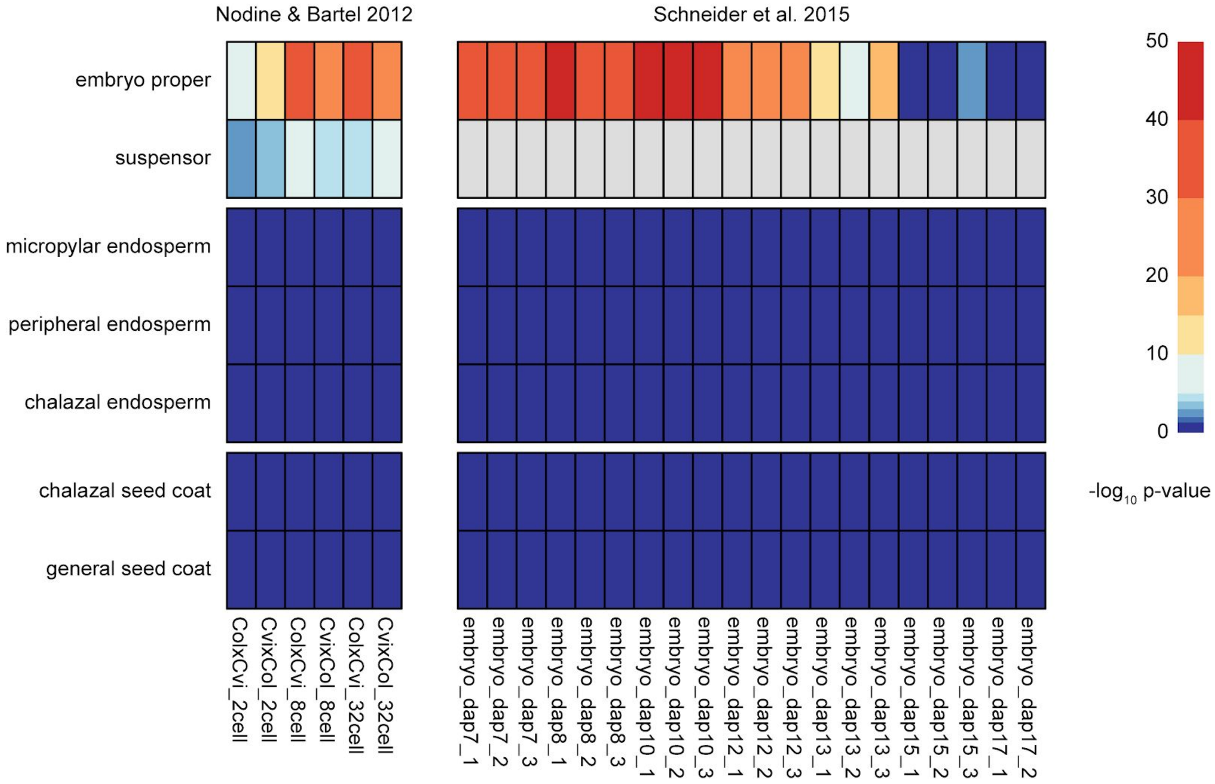


**Figure S1.** Tissue-enrichment-test for referenced embryo transcriptomes

Heatmap of the p-values obtained from the *tissue-enrichment test* (Schon and Nodine 2017). We performed the test on the early (*left*, (Nodine and Bartel 2012)) and late embryo samples (*right*, (Schneider et al. 2016)). Early embryo data (Nodine and Bartel 2012) was reanalyzed as quantification was performed differently from (Schon and Nodine 2017) (see Materials & Methods for details).

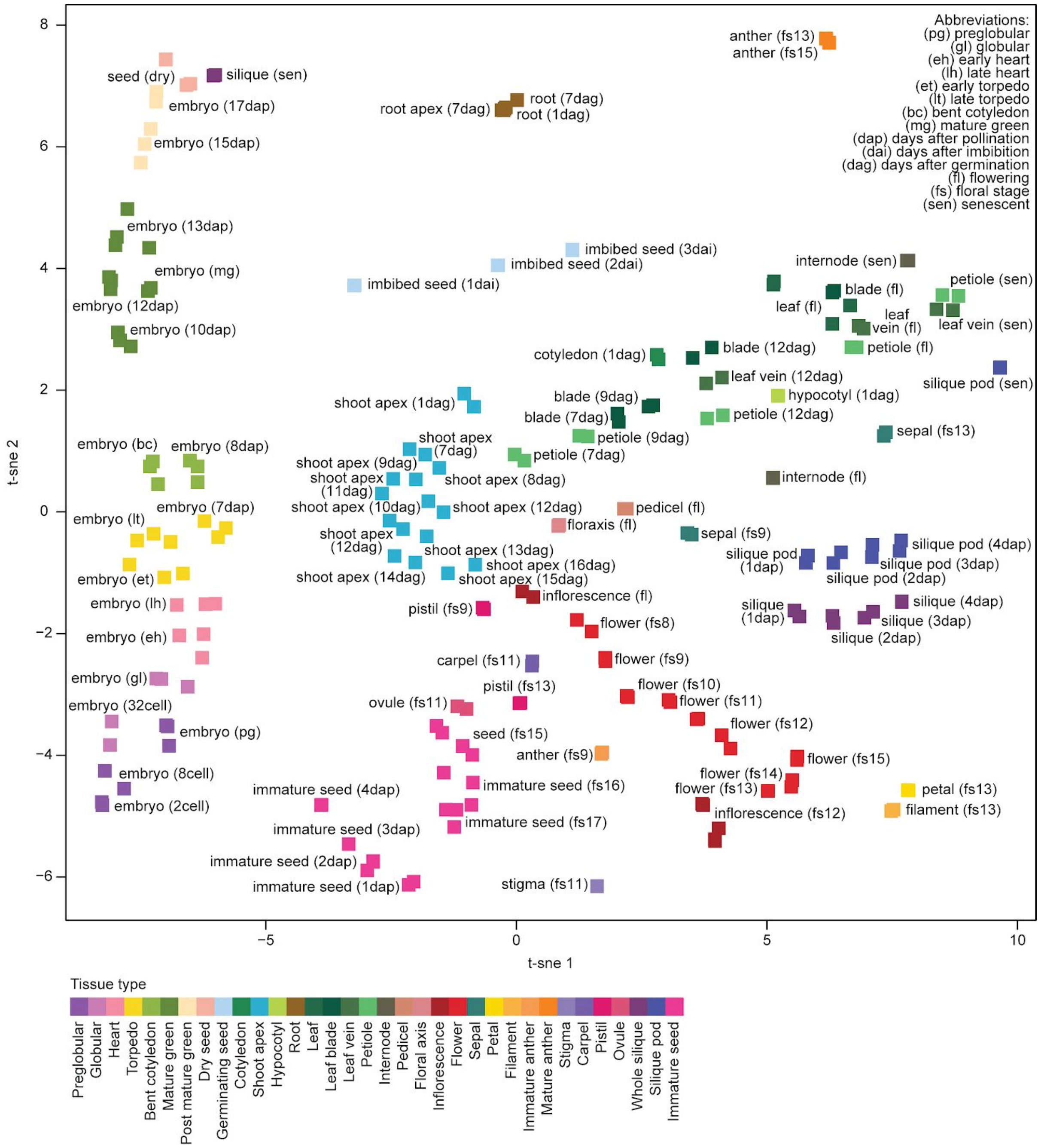

**Figure S2.** t-SNE of all samples

T-SNE comparing transcriptomes from different tissues and sources (Nodine and Bartel 2012; Belmonte et al. 2013; Klepikova et al. 2015, 2016; Schneider et al. 2016; Lutzmayer et al. 2017; Schon et al. 2018). Every square represents one sample (see also table S 1). Samples are colored by tissue type and labeled with specific developmental age.

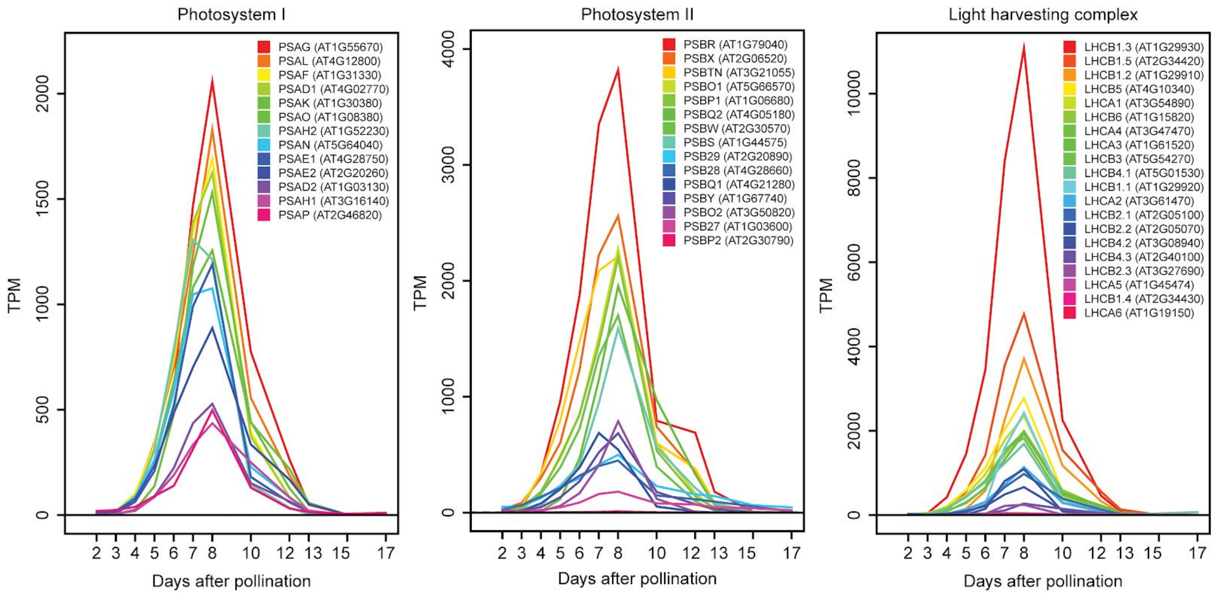


**Figure S3.** Expression of photosynthesis pathway components during embryogenesis

Gene expression (TPM) of the annotated, nuclear encoded components of photosystem I (*left*), photosystem II (*center*) and the light-harvesting complexes 1 & 2 (*right*).

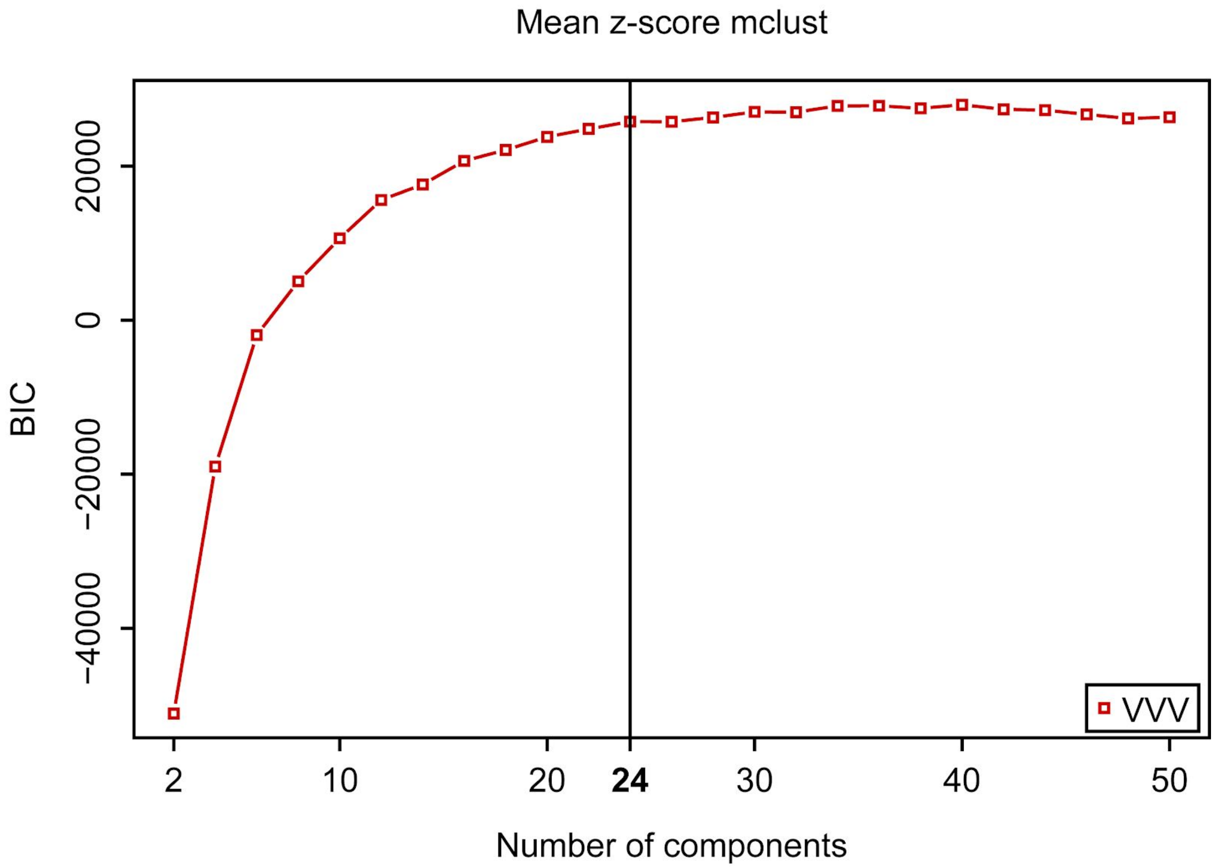


**Figure S4.** BIC score for model based clustering

Mclust (Scrucca et al. 2016) calculated BIC score visualized against the different numbers of clusters. The clustering was performed via the *VVV* (ellipsoidal, varying volume, shape, and orientation) model. The first local maximum BIC is marked with a vertical line (see Materials & Methods).

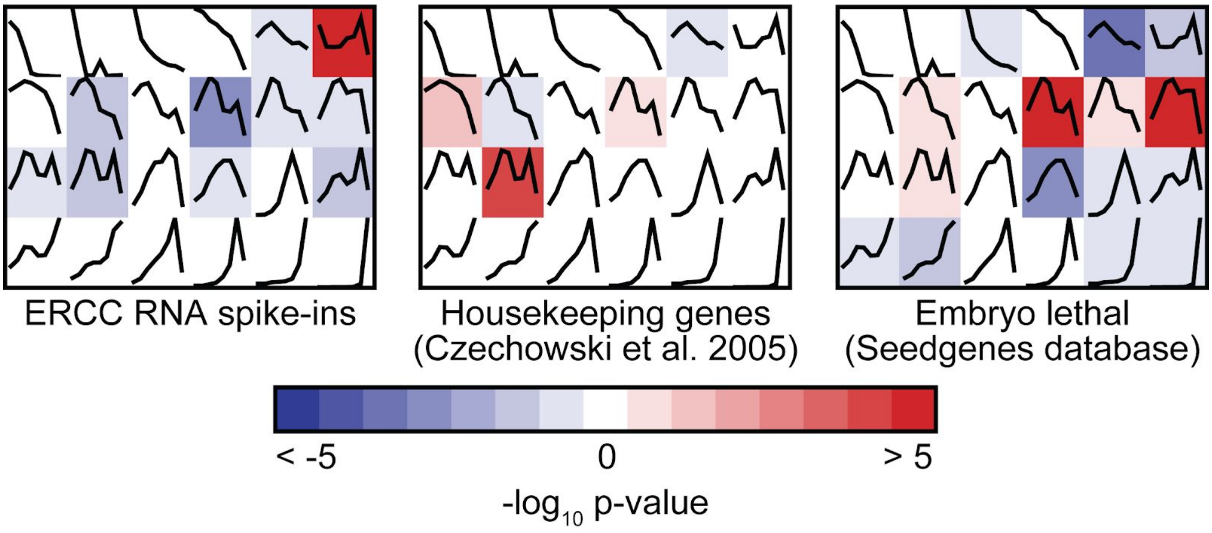


**Figure S5.** Gene group enrichment for model based clustering

Overview which clusters are either significantly enriched (*red*) or depleted (*blue*) for ERCC RNA spike-ins (Baker et al. 2005) (*left*) housekeeping genes (Czechowski et al. 2005) (*mid*), or embryo lethal genes (Meinke et al. 2008) (*right*). P-value: hypergeometric distribution, Benjamini-Hochberg multiple testing correction.

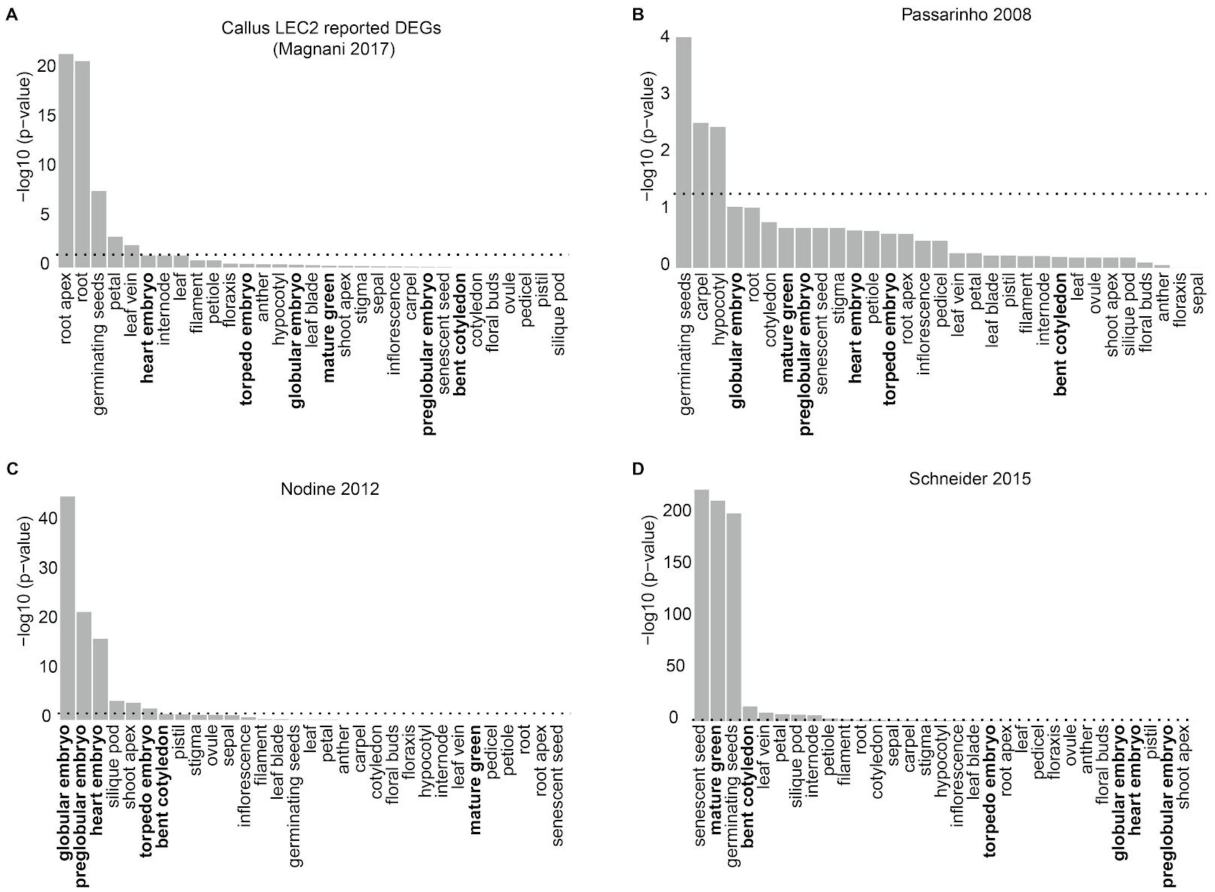


**Figure S6.** Additional TissueEnrich results

Tissue-specific gene enrichment (Jain and Tuteja 2018) of the DEGs originally reported by Magnani and colleagues (Magnani et al. 2017) **(A),** of a BBM overexpression line that forms somatic embryos on leafs (Passarinho et al. 2008) **(B)**, an early (Nodine and Bartel 2012) **(C)** and a late embryo (Schneider et al. 2016) **(D)** time series. All DEGs where subsetted for genes expressed over 1 TPM and having a VST fold change of at least 2, except for **(B)**, which was obtained from microarray data. Tissue enrichment testing was perfomed against (Klepikova et al. 2015, 2016; Lutzmayer et al. 2017; Schon et al. 2018).

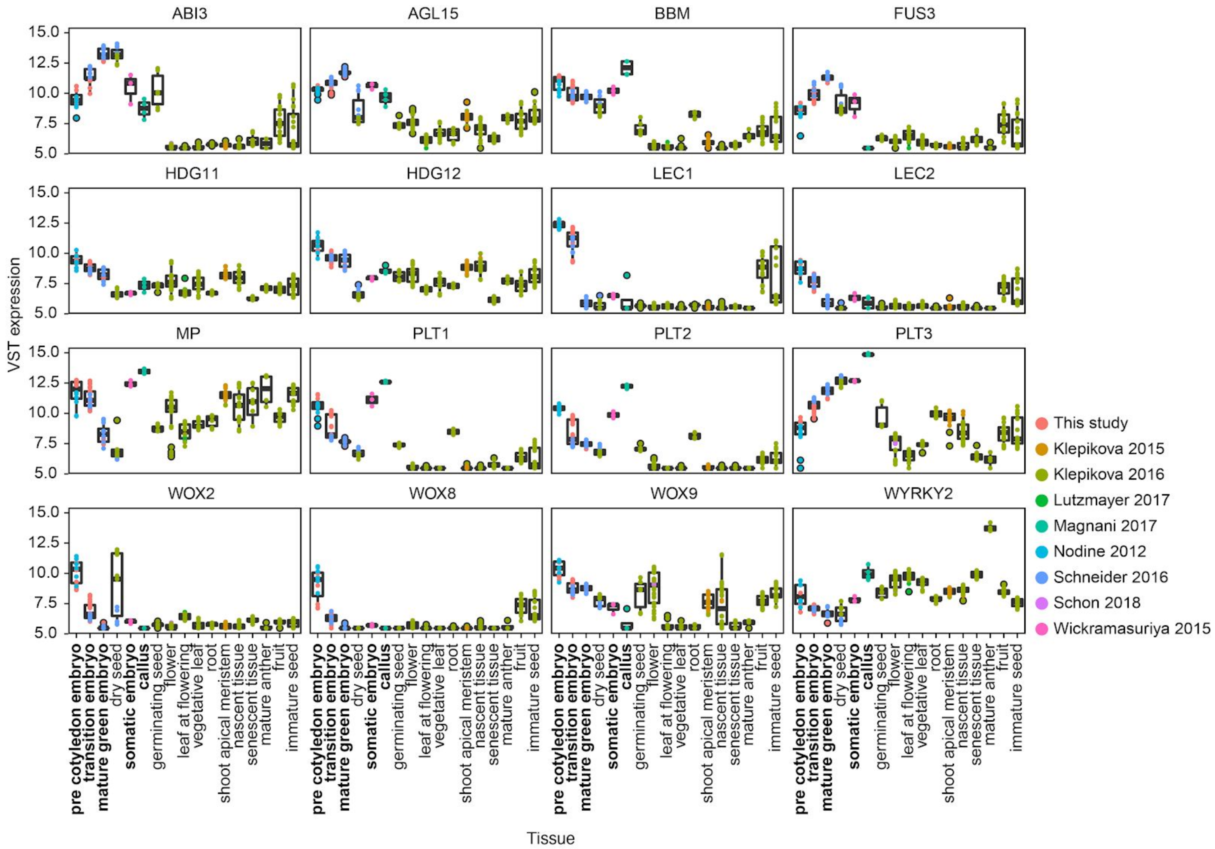


**Figure S7.** Expression of select embryonic markers genes

VST normalized expression (Love et al. 2014) of several select embryonic markers (Lau et al. 2012) in in their respective tissue cluster (see table S1 & Fig S2). The data source is indicated by the respective color (Nodine and Bartel 2012; Wickramasuriya and Dunwell 2015; Klepikova et al. 2015, 2016; Schneider et al. 2016; Lutzmayer et al. 2017; Magnani et al. 2017; Schon et al. 2018).

**Online Tables**

**Table S1.** Sample overview

- Can be found under filename Table_S1_sample_table.xlsx
- Contains metadata on all RNA-seq samples generated or analyzed in this study

**Table S2.** Expression values

- Can be found under filename Table_S2_expression_values.xlsx
- Contains the different types of normalized expression values used in this study

**Table S3.** Pearson correlation matrix

- Can be found under filename Table_S3_correlation_matrix.xlsx
- Contains a matrix with pairwise Pearson correlation coefficients of all samples (on $\log_2$(TPM+1) data.

**Table S4.** Tissue-cluster enriched genes

- Can be found under filename Table_S4_cluster_enriched_genes.xlsx
- Contains ANOVA enrichment results for all tissue clusters

**Table S5.** Tissue-cluster depleted genes

- Can be found under filename Table_S5_cluster_depleted_genes.xlsx
- Contains ANOVA depletion results for all tissue clusters

**Table S6.** Model based clustering

- Can be found under filename Table_S6_mclust.xlsx
- Contains covariance cluster information for all genes and Gene Ontology term enrichments for each cluster

**Table S7.** Phase and stage specific markers

- Can be found under the filename Table_S7_markers.xlsx
- Contains the MGFR (El Amrani et al. 2015) identified marker genes for the different embryonic phases as well as for all embryonic stages

**Table S8.** DEGs and TissueEnrich results

- Can be found under the filename Table_S8_enrichments.xlsx
- Contains the identified DEGs from the somatic embryogenesis part, as well as the results from the *TissueEnrich* tests (Jain and Tuteja 2018) and sample names we tested against (see Table S1 & S2 for more info on the samples)

**Online References**


Baker SC, Bauer SR, Beyer RP, et al (2005) The External RNA Controls Consortium: a progress report. Nat Methods 2:731–734. doi: 10.1038/nmeth1005-731

Belmonte MF, Kirkbride RC, Stone SL, et al (2013) Comprehensive developmental profiles of gene activity in regions and subregions of the Arabidopsis seed. Proc Natl Acad Sci U S A 110:E435–44. doi: 10.1073/pnas.1222061110

Czechowski T, Stitt M, Altmann T, et al (2005) Genome-wide identification and testing of superior reference genes for transcript normalization in Arabidopsis. Plant Physiol 139:5–17. doi: 10.1104/pp.105.063743

El Amrani K, Stachelscheid H, Lekschas F, et al (2015) MGFM: a novel tool for detection of tissue and cell specific marker genes from microarray gene expression data. BMC Genomics 16:645. doi: 10.1186/s12864-015-1785-9

Jain A, Tuteja G (2018) TissueEnrich: Tissue-specific gene enrichment analysis. Bioinformatics. doi: 10.1093/bioinformatics/bty890

Klepikova AV, Kasianov AS, Gerasimov ES, et al (2016) A high resolution map of the Arabidopsis thaliana developmental transcriptome based on RNA-seq profiling. Plant J 88:1058–1070. doi: 10.1111/tpj.13312

Klepikova AV, Logacheva MD, Dmitriev SE, Penin AA (2015) RNA-seq analysis of an apical meristem time series reveals a critical point in Arabidopsis thaliana flower initiation. BMC Genomics 16:466. doi: 10.1186/s12864-015-1688-9

Lau S, Slane D, Herud O, et al (2012) Early embryogenesis in flowering plants: setting up the basic body pattern. Annu Rev Plant Biol 63:483–506. doi: 10.1146/annurev-arplant-042811-105507

Love MI, Huber W, Anders S (2014) Moderated estimation of fold change and dispersion for RNA-seq data with DESeq2. Genome Biol 15:550. doi: 10.1186/s13059-014-0550-8

Lutzmayer S, Enugutti B, Nodine MD (2017) Novel small RNA spike-in oligonucleotides enable absolute normalization of small RNA-Seq data. Sci Rep 7:5913. doi: 10.1038/s41598-017-06174-3

Magnani E, Jiménez-Gómez JM, Soubigou-Taconnat L, et al (2017) Profiling the onset of somatic embryogenesis in Arabidopsis. BMC Genomics 18:998. doi: 10.1186/s12864-017-4391-1

Meinke D, Muralla R, Sweeney C, Dickerman A (2008) Identifying essential genes in Arabidopsis thaliana. Trends Plant Sci 13:483–491. doi: 10.1016/j.tplants.2008.06.003

Nodine MD, Bartel DP (2012) Maternal and paternal genomes contribute equally to the transcriptome of early plant embryos. Nature 482:94–97. doi: 10.1038/nature10756

Passarinho P, Ketelaar T, Xing M, et al (2008) BABY BOOM target genes provide diverse entry points into cell proliferation and cell growth pathways. Plant Mol Biol 68:225–237. doi: 10.1007/s11103-008-9364-y

Schneider A, Aghamirzaie D, Elmarakeby H, et al (2016) Potential targets of VIVIPAROUS1/ABI3-LIKE1 (VAL1) repression in developing Arabidopsis thaliana embryos.

Plant J 85:305–319. doi: 10.1111/tpj.13106

Schon MA, Kellner MJ, Plotnikova A, et al (2018) NanoPARE: parallel analysis of RNA 5' ends from low-input RNA. Genome Res. doi: 10.1101/gr.239202.118

Schon MA, Nodine MD (2017) Widespread Contamination of Arabidopsis Embryo and Endosperm Transcriptome Data Sets. Plant Cell 29:608–617. doi: 10.1105/tpc.16.00845

Scrucca L, Fop M, Murphy TB, Raftery AE (2016) mclust 5: Clustering, Classification and Density Estimation Using Gaussian Finite Mixture Models. R J 8:289–317

Soneson C, Love MI, Robinson MD (2015) Differential analyses for RNA-seq: transcript-level estimates improve gene-level inferences. F1000Res 4:1521. doi: 10.12688/f1000research.7563.2

Wickramasuriya AM, Dunwell JM (2015) Global scale transcriptome analysis of Arabidopsis embryogenesis in vitro. BMC Genomics 16:301. doi: 10.1186/s12864-015-1504-6
